# Localized Chromatin Rewiring Shapes Rhythmic Transcription within a Stable 3D Genome in Rice

**DOI:** 10.64898/2026.10.23.761304

**Authors:** Ying Zhang, Guoting Chen, Liangliang Guo, Mengxing Cao, Jixian Zhou, Xingwang Li, Jiapei Yan

## Abstract

How spatial genome architecture coordinates rhythmic gene transcription across scales in plants remains poorly understood. Here, we integrated bridge linker Hi-C (BL-Hi-C), H3K9ac ChIP-seq and RNA-seq at six time points across a 24-h circadian cycle in rice. While global higher-order structure was largely invariant, with 84.5% of A/B compartments and over 95% of topologically associating domains (TADs) conserved across all time points, we identified 557 dynamic compartment transitions coupled to changes in active chromatin marks. Within this stable structural scaffold, 35.8% of TADs containing clustered loci of multiple rhythmic genes with synchronized expression phases. In contrast, fine-scale intradomain contacts and chromatin loops exhibited pronounced temporal plasticity. Chromatin interactions connected rhythmic genes with diverse expression phases. Myb-binding motifs were enriched at chromatin interaction anchors at dawn and early day (ZT0 and ZT4). Together, our findings demonstrate that localized chromatin rewiring occurs within relatively stable architectural frameworks, providing a spatiotemporal context for circadian gene regulation in rice.

## Introduction

The circadian clock coordinates daily transcriptional programs, photoperiodic flowering and metabolic processes in rice. Clock-dependent regulation determines the critical day length for flowering, linking endogenous rhythms to photoperiod responses [1]. Under field conditions, disruption of clock function alters the phase and amplitude of widespread transcriptional rhythms and changes phenylpropanoid metabolism [2]. These findings establish circadian regulation as an important component of rice reproductive development and metabolic organization. The breadth of rhythmic transcription also raises questions about its organization across the genome. Examining chromatin architecture throughout the circadian cycle can therefore clarify the spatial context in which these transcriptional programs operate.

Chromosome conformation capture methods resolve genome organization at several scales. Hi-C identified broad compartments associated with distinct chromatin properties [3] and topologically associating domains (TADs) with relatively frequent internal contacts [4]. Rice genomes likewise exhibit compartmentalization, local interaction domains and chromatin loops associated with genomic and epigenetic features [5]. Compartment identity describes large-scale segregation of chromatin regions, whereas domain boundaries and individual contacts describe local insulation and interactions between specific regions, respectively. These levels may differ in their temporal stability, allowing transcription-associated contact changes within a persistent higher-order framework.

Mammalian studies demonstrate links between daily transcriptional programs and chromosome architecture. In mouse liver, REV-ERBα regulates enhancer–promoter contacts through a corepressor-associated mechanism involving histone deacetylation [6]. Perturbation of a contacted regulatory element at a clock locus has also linked chromatin topology to transcriptional phase and behavior [7]. Genome-wide and promoter-capture Hi-C analyses identified time-dependent compartment assignments and promoter contacts alongside relatively stable domain boundaries [8]. These findings motivate analysis across structural scales while leaving open how temporal genome organization operates in plants.

In rice, RNAPII-associated ChIA-PET revealed time-dependent interaction networks linked to rhythmic gene expression [9]. Integrated epigenomic and transcriptomic profiling further identified associations of transcriptional rhythms with H3K9ac and transcription factor occupancy, while BL-Hi-C comparisons at ZT8 and ZT20 under constant light identified temporal variation in chromatin interactions [10]. These studies establish a basis for investigating gene-associated contact dynamics within genome-wide architecture throughout the sampled cycle. Profiling additional time points can distinguish compartment transitions, domain stability and local interaction dynamics, and assess their associations with the expression phases of interacting genes.

Here, we combined bridge linker Hi-C (BL-Hi-C) [11], H3K9ac ChIP-seq and RNA-seq at ZT0, ZT4, ZT8, ZT12, ZT16 and ZT20 to characterize temporal genome organization and its association with rhythmic transcription in rice. We assessed the temporal stability of compartments and domains, the chromatin features associated with compartment switching, and contact variation within domains containing rhythmic genes. We then examined the temporal dynamics of chromatin interactions involving rhythmic genes and their associations with expression phases, histone acetylation and enriched regulatory motifs. Together, these analyses reveal selective compartment transitions and dynamic interactions involving rhythmic genes within a largely conserved higher-order framework.

## Results

### Genome-Wide Profiling Identifies Oscillating Chromatin Compartments in Rice

To map temporal genome organization in rice, we integrated BL-Hi-C, H3K9ac ChIP-seq and RNA-seq profiles from wild-type seedlings sampled at ZT0, ZT4, ZT8, ZT12, ZT16 and ZT20 over 24 h (Fig. 1a). The BL-Hi-C datasets yielded approximately 100 million uniquely mapped paired-end tags (PETs). First principal component (PC1) eigenvectors from 100-kb interaction matrices defined A and B compartments across the 12 rice chromosomes (Fig. 2a). Of the 3,602 regions analyzed, 3,045 (84.5%) retained the same compartment identity, whereas 557 (15.5%) switched identity during the sampled cycle (Fig. 1b). We operationally designated these switching regions oscillating chromatin compartments (OCCs).

**Fig. 1.**
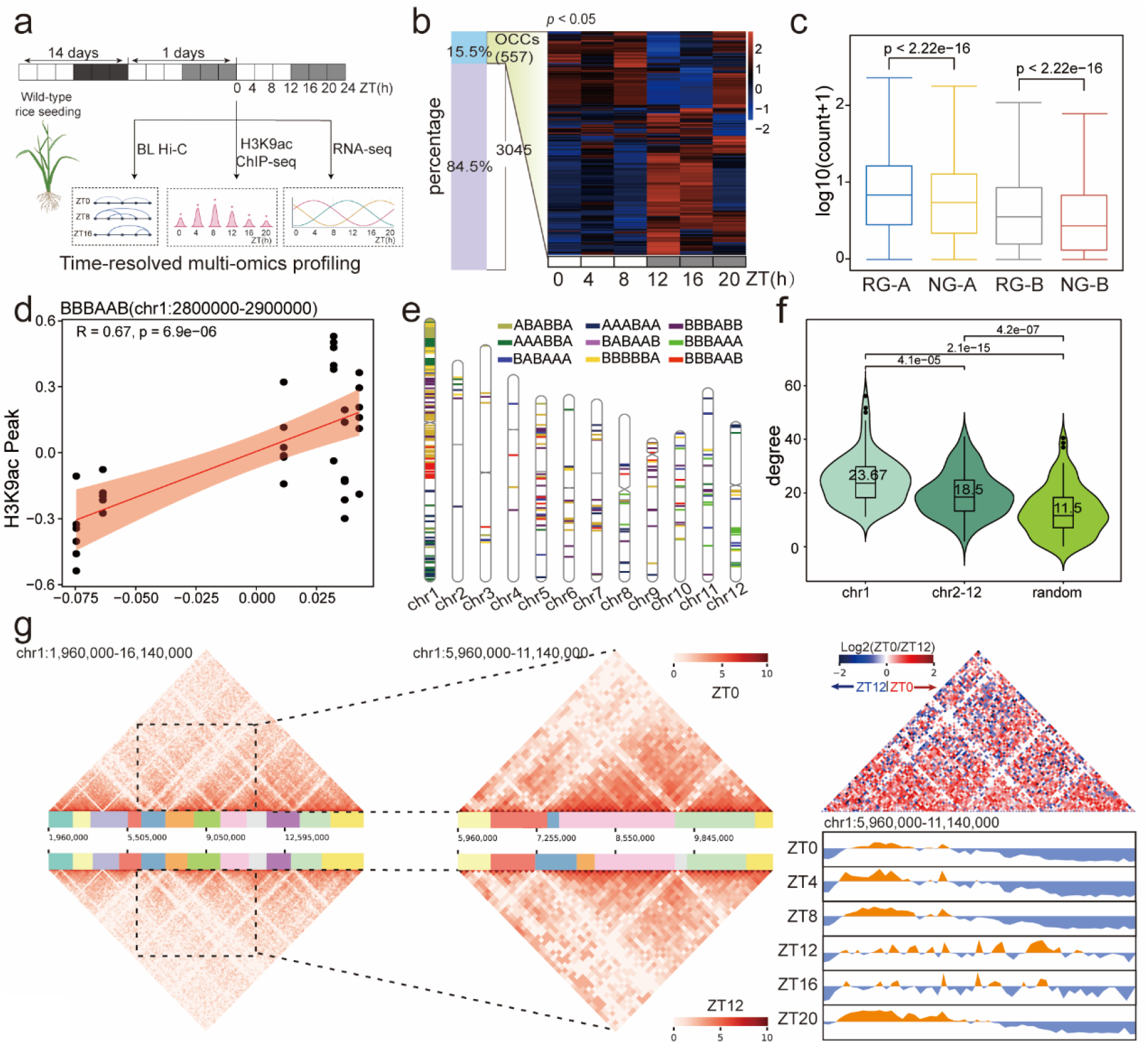
Time-dependent switching of chromatin compartments in rice. a, Experimental design for time-resolved profiling of wild-type rice seedlings. BL-Hi-C, H3K9ac ChIP-seq and RNA-seq profiles were analyzed at ZT0, ZT4, ZT8, ZT12, ZT16 and ZT20 over a 24-h cycle. b, Classification of genomic regions by temporal compartment identity. Left, proportions of OCCs (15.5%, n = 557) and stable regions (84.5%, n = 3,045). Right, heatmap of PC1-derived compartment profiles for OCCs across the six time points. OCCs change A/B assignment during the sampled cycle; stable regions retain the same assignment. **c**, Expression levels of rhythmic and non-rhythmic genes within OCCs, grouped by compartment state and represented as log₁₀(count + 1). RG-A and NG-A denote rhythmic and non-rhythmic genes in A compartments; RG-B and NG-B denote the corresponding groups in B compartments. Brackets indicate comparisons between rhythmic and non-rhythmic genes within each compartment state. d, Relationship between compartment score and H3K9ac peak signal at a representative BBBAAB locus on chr1 (2,800,000–2,900,000 bp; reported R = 0.67, P = 6.9 × 10⁻⁶). The red line shows the fitted linear relationship, and the shaded band indicates uncertainty around the fit. **e**, Chromosomal distribution of nine abundant OCC switching patterns. Colors distinguish the patterns indicated in the legend. Each six-letter code specifies compartment identity at ZT0, ZT4, ZT8, ZT12, ZT16 and ZT20, in that order. f, Interaction degree of rhythmic genes on chr1, rhythmic genes on chr2–12 and a random gene comparison set. Summary values shown in the plots are 23.67, 18.5 and 11.5, respectively. Brackets indicate the pairwise comparisons, with P values above. **g**, Representative BL-Hi-C contact maps of chr1:1.96–16.14 Mb at ZT0 and ZT12. Dashed boxes identify the expanded interval, chr1:5.96–11.14 Mb. Right, differential interaction map, expressed as log₂(ZT0/ZT12), and PC1 eigenvector tracks across the six time points. Red and blue in the differential map indicate relatively higher interaction signals at ZT0 and ZT12, respectively. ZT, Zeitgeber time; PC1, first principal component; RNAPII, RNA polymerase II.

**Fig. 2.**
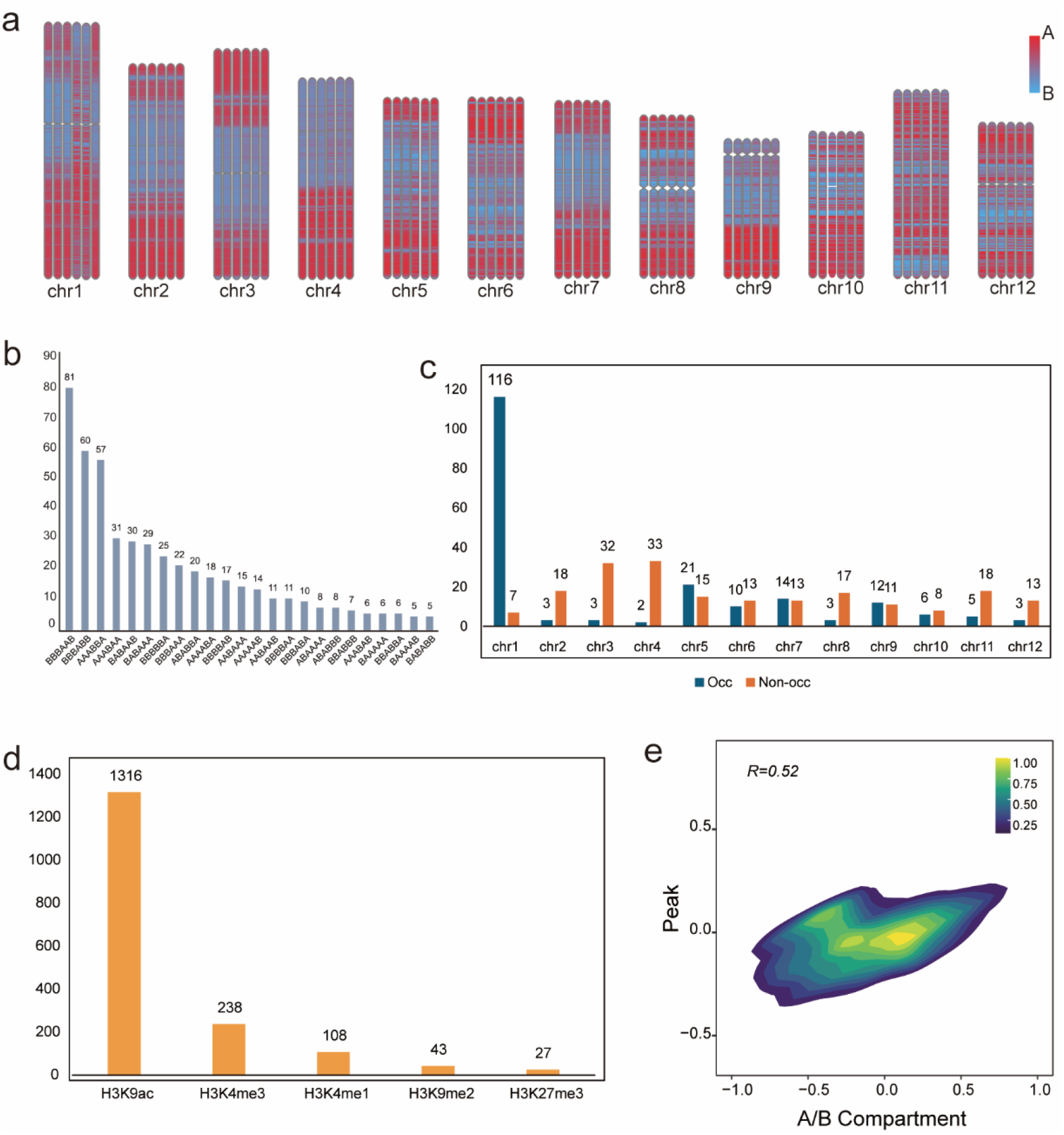
Genome-wide compartment profiles and characteristics of switching regions. a, A/B compartment profiles across the 12 rice chromosomes at six time points. Red and blue indicate A and B assignments, respectively. b, Counts of OCC regions by switching pattern. Each code lists compartment states at ZT0, ZT4, ZT8, ZT12, ZT16 and ZT20, in that order. c, Chromosomal distributions of the selected OCC (blue) and non-OCC (orange) comparison sets. d, Counts of histone-mark features included in the compartment-related analysis: H3K9ac, 1,316; H3K4me3, 238; H3K4me1, 108; H3K9me2, 43; and H3K27me3, 27. e, Compartment score versus peak signal (reported R = 0.52); color indicates observation density.

Compartment switching was unevenly distributed across the genome. The most abundant patterns included BBBAAB, BBBABB and AAABBA, with each code specifying compartment identity in chronological sampling order (Fig. 2b). The nine abundant patterns showed a pronounced concentration on chr1 (Fig. 1e). In the selected comparison sets, chr1 contained 116 of 198 OCC regions and seven non-OCC regions (Fig. 2c).

Rhythmic genes on chr1 also had higher interaction degree than those on chr2-12 and the random comparison set (P = 4.1 × 10⁻⁵ and 2.1 × 10⁻¹⁵, respectively; Fig. 1f). Regional contact maps illustrated the structural changes associated with this compartment variation. Maps of chr1:1.96-16.14 Mb at ZT0 and ZT12, together with an expanded differential map of chr1:5.96-11.14 Mb, showed local changes in contact patterns. PC1 tracks across all six time points showed changes in compartment identity within the same region (Fig. 1g).

### Dynamic Compartment Switching Couples with Rhythmic Histone Acetylation

To characterize chromatin features associated with compartment switching, we examined temporal profiles of histone modifications and accessibility. H3K9ac and H3K27ac had the highest proportions of rhythmic features: 60.9% (11,750 of 19,285) and 53.7% (9,481 of 17,649), respectively. By comparison, 153 H3K4me3 and four H3K4me1 features were classified as rhythmic, whereas no rhythmic H3K27me3, H3K9me2 or ATAC-seq features were detected under the applied criteria (Fig. 3a).

**Fig. 3.**
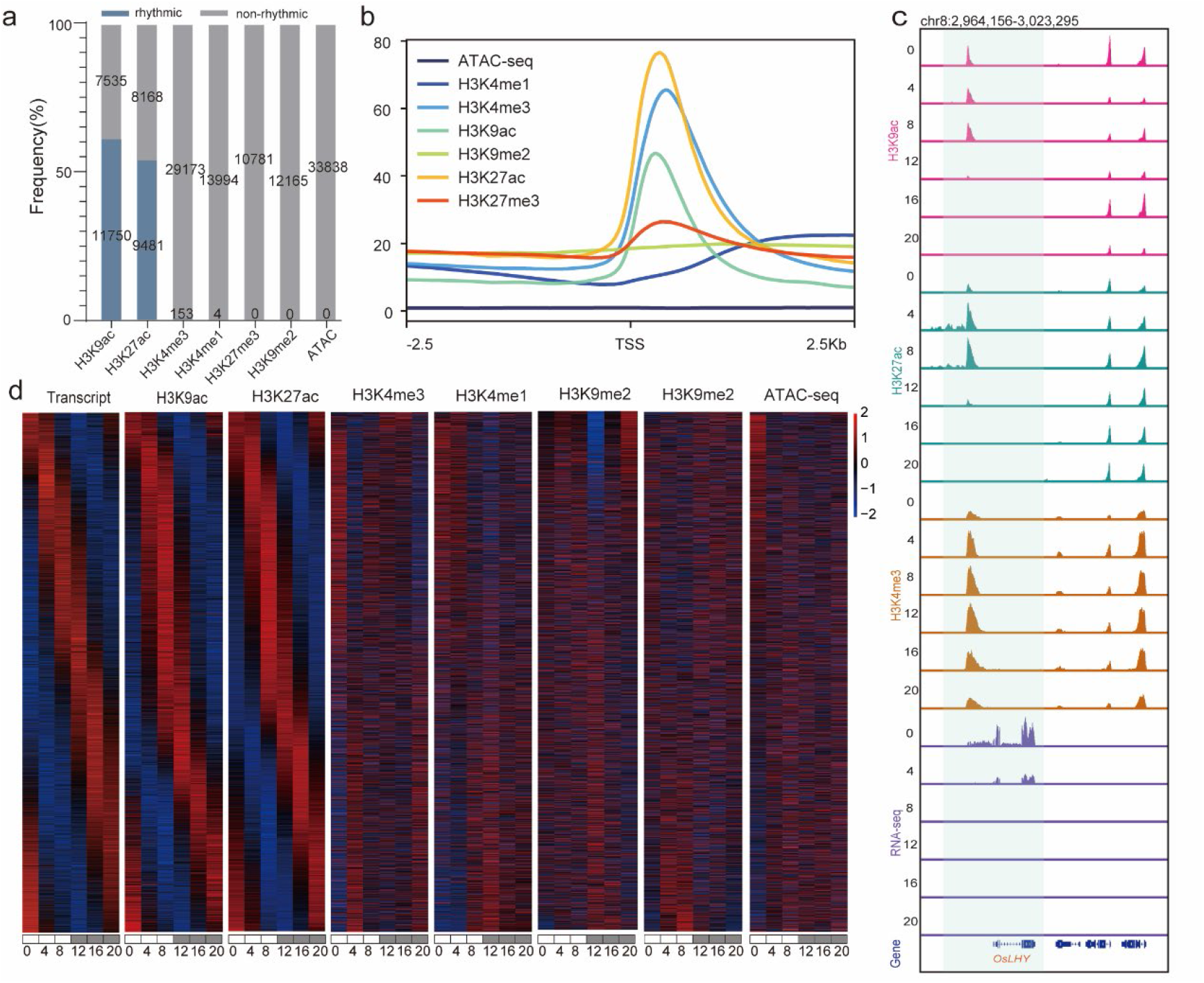
Temporal profiles of histone modifications, chromatin accessibility and transcription. a, Proportions of features classified as rhythmic or non-rhythmic in the indicated histone-modification and ATAC-seq datasets. Blue and gray indicate rhythmic and non-rhythmic features, respectively. Numbers indicate feature counts. An absence of detected rhythmic features refers to the applied analysis criteria. b, Average ATAC-seq and histone-modification signal profiles within 2.5 kb upstream and downstream of transcription start sites (TSSs). c, Genome-browser tracks for H3K9ac, H3K27ac, H3K4me3 and RNA-seq at ZT0, ZT4, ZT8, ZT12, ZT16 and ZT20 across chr8:2,964,156–3,023,295, encompassing *OsLHY*. Shading highlights the *OsLHY* region; gene models are shown below. d, Heatmaps of transcript abundance and histone-modification and ATAC-seq signals across six time points. Red and blue indicate relatively high and low values on the displayed scale, respectively.

H3K9ac, H3K27ac and H3K4me3 were enriched near transcription start sites (Fig. 3b). Browser tracks at *OsLHY* and genome-wide heatmaps further illustrated temporal variation in transcription and chromatin signals (Fig. 3c, d).

We then examined associations of transcription and chromatin signals with compartment state. Within OCCs, rhythmic genes showed higher expression than non-rhythmic genes in both A and B compartments (P < 2.22 × 10⁻¹⁶ for each comparison; Fig. 1c). The compartment-associated histone-mark analysis included 1,316 H3K9ac, 238 H3K4me3, 108 H3K4me1, 43 H3K9me2 and 27 H3K27me3 features (Fig. 2d). At a representative BBBAAB locus on chr1 (2.80–2.90 Mb), compartment score was positively associated with H3K9ac peak signal (reported R = 0.67, P = 6.9 × 10⁻⁶; Fig. 1d). A separate compartment-score–peak-signal comparison also showed a positive association (reported R = 0.52; Fig. 2e).

These associations were also evident in the temporal chromatin profiles of the three abundant switching patterns. In BBBAAB regions, RNAPII occupancy and H3K9ac enrichment were generally higher at ZT12 and ZT16, when the regions occupied A compartments, whereas H3K9me2 enrichment was lower. AAABBA regions showed higher RNAPII and H3K9ac signals during their A-compartment time points (ZT0, ZT4, ZT8 and ZT20), while BBBABB regions showed the corresponding pattern at ZT12 (Fig. 4a). Together, these observations associate selective compartment transitions with changes in transcription-related chromatin signals within a predominantly stable compartment landscape.

**Fig. 4.**
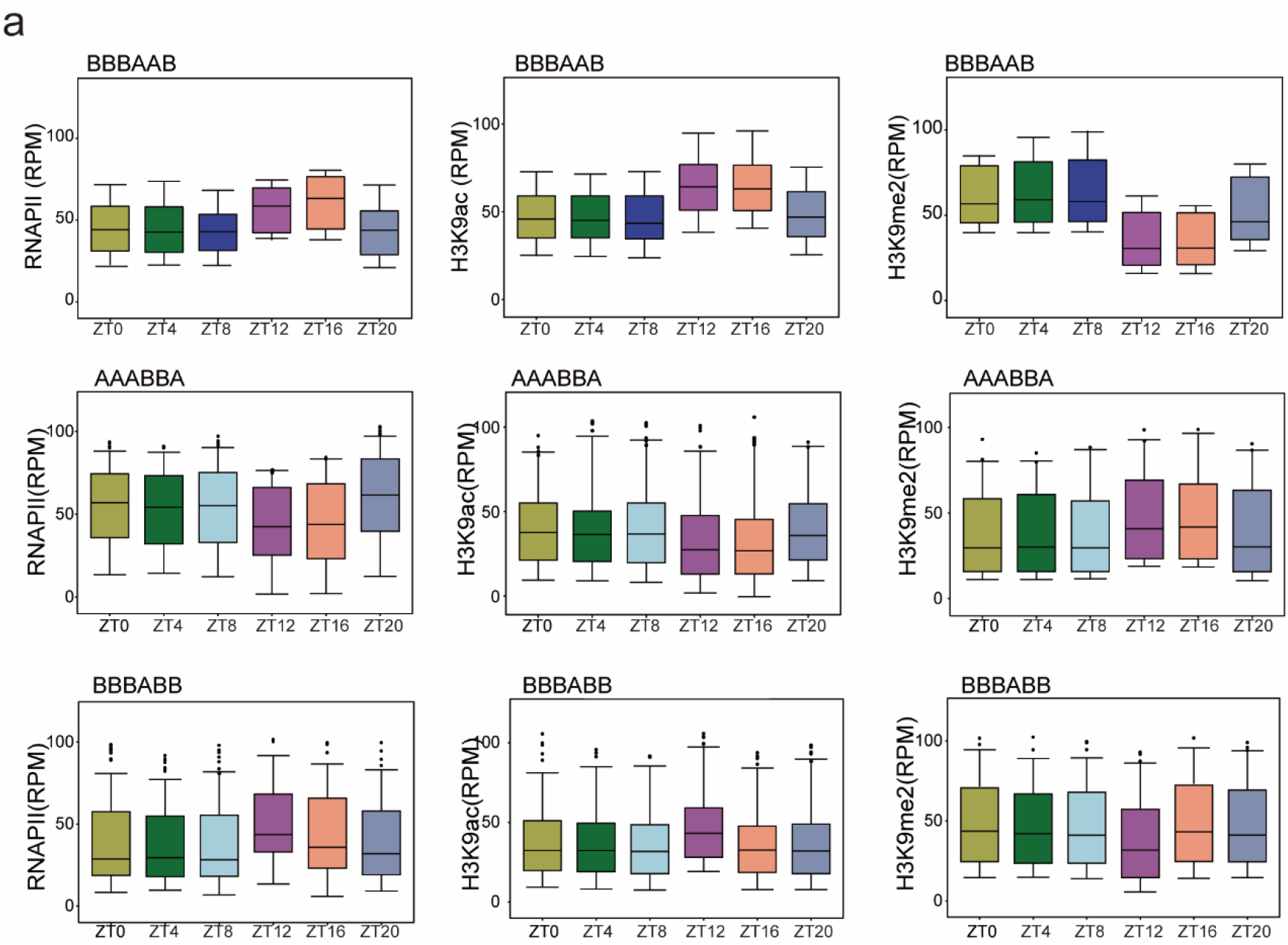
Temporal RNAPII and histone-modification profiles of compartment-switching regions. a, RNAPII, H3K9ac and H3K9me2 signal distributions across six time points for BBBAAB (top), AAABBA (middle) and BBBABB (bottom) OCCs. Signals are expressed as reads per million (RPM). BBBAAB regions occupy A compartments at ZT12 and ZT16, AAABBA regions at ZT0, ZT4, ZT8 and ZT20, and BBBABB regions at ZT12. Each pattern occupies B compartments at the remaining time points. Letters in each code follow chronological sampling order.

### Stable TAD Organization Accommodates Temporal Variation in Internal Contacts

In contrast to the dynamic behavior of OCCs, domain-level organization of the rice genome remained largely stable over the 24-h cycle. Topologically associating domain (TAD) calling identified 1,078 domains shared across all six time points, representing more than 95% of all identified TADs at any given time point (Fig. 5e). This demonstrates that the architectural boundaries delimiting chromatin domains are largely refractory to diurnal fluctuations.

**Fig. 5.**
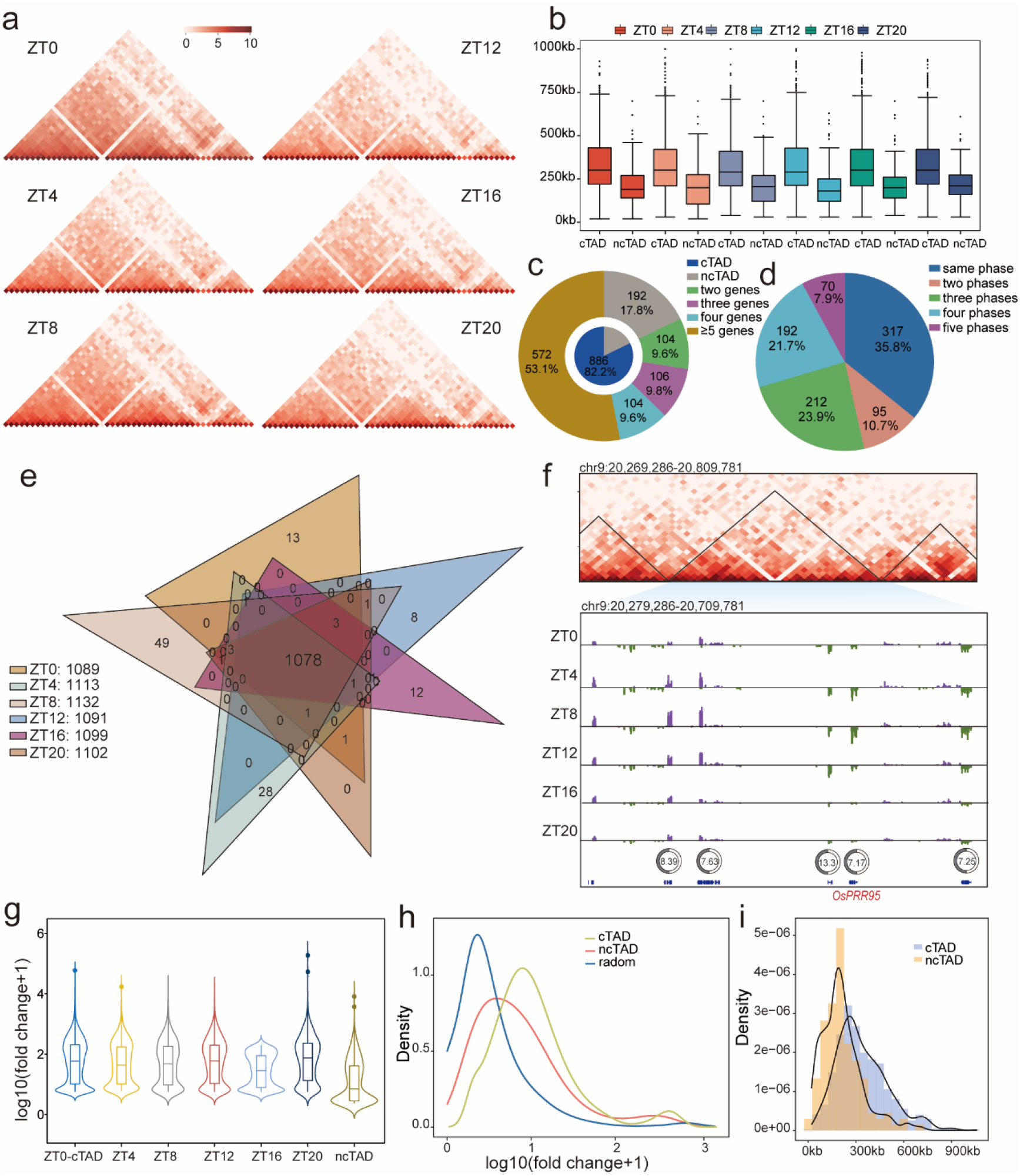
Conserved domain organization and temporal variation in internal chromatin contacts. a, Aggregate BL-Hi-C contact maps for 100 randomly selected TADs identified at ZT0, shown across six time points (ZT0–ZT20). The maps summarize temporal similarity in domain organization. b, Genomic size distributions of cTADs and ncTADs at each time point. cTADs contain at least two rhythmic genes; ncTADs contain fewer than two. c, Nested pie chart showing the proportions of cTADs and ncTADs (inner circle) and domains grouped by rhythmic-gene number, including the ncTAD category (outer ring). The analyzed set comprises 886 cTADs and 192 ncTADs; percentages are relative to all 1,078 domains. d, Distribution of expression-phase categories among the 886 cTADs. In 317 cTADs (35.8%), rhythmic genes share the same phase category; the remaining sectors indicate domains containing genes assigned to two, three, four or five phase categories. e, Overlap of TADs identified at the six time points. The central intersection contains 1,078 domains detected at every time point. f, Representative region containing *OsPRR95*. Top, BL-Hi-C contact map and domain boundaries across chr9:20,269,286–20,809,781. Bottom, RNA-seq tracks at the six time points and gene models across chr9:20,279,286–20,709,781. Clock symbols indicate estimated expression phases in hours. g, Distributions of the intradomain interaction fold-change metric, expressed as log₁₀(fold change + 1), for cTADs in each of the six time-point groups and for the ncTAD comparison group. h, Density distributions of the transformed intradomain interaction fold-change metric for cTADs, ncTADs and the random TAD comparison set. i, Histograms and density curves showing the genomic size distributions of cTADs and ncTADs.

Within this largely conserved structural framework, we examined whether TADs spatially group rhythmic genes. We first designated domains containing at least two rhythmic genes as circadian TADs (cTADs) and the remaining domains as non-circadian TADs (ncTADs). These operational labels describe gene content rather than domain rhythmicity. The analyzed set comprised 886 cTADs (82.2%) and 192 ncTADs (17.8%); 104, 106, 104 and 572 cTADs contained two, three, four and at least five rhythmic genes, respectively (Fig. 5c). cTADs generally spanned larger genomic intervals than ncTADs at each time point, and their size distribution extended toward longer domains (Fig. 5b, i).

We next examined whether cTADs spatially group rhythmic genes with shared expression phases. In 317 cTADs (35.8%), rhythmic genes were assigned to the same expression-phase category (Fig. 5d), identifying a subset of domains containing genes that peak within a shared temporal window. The remaining cTADs contained two (95, 10.7%), three (212, 23.9%), four (192, 21.7%) or five (70, 7.9%) phase categories.

For example, the TAD containing the core clock gene *OsPRR95* comprised five rhythmic genes. *OsPRR95* had an estimated expression phase of 7.17 h, and the three of the other labelled genes peaked at approximately 7 h (Fig. 5f). These observations suggest that TADs can provide a spatial context for coordinated peak expression of subsets of rhythmic genes during the sampled cycle.

To assess contact variation within these conserved domains, we compared intradomain interactions across the sampled cycle. The intradomain interaction fold-change distributions, expressed as log₁₀(fold change + 1), overlapped across the six cTAD time-point groups but differed in their central values and upper tails (Fig. 4g). The principal peak of the cTAD distribution occurred at a higher value than those of ncTADs and the random TAD comparison set (Fig. 5h). These distributions indicate that contacts varied within the largely conserved domain framework.

Together, these results suggest that while TAD boundaries remain static across the day and night, they function as insulated regulatory neighborhoods that spatially organize co-expressed, phase-synchronized rhythmic genes.

### Temporal Rewiring of Chromatin Loops Connects Synchronous Rhythmic Genes

Despite the overarching stability of TAD boundaries, fine-scale chromatin contacts within domains exhibited substantial temporal plasticity. Rhythmic anchor genes accounted for 22.9–27.1% of interacting partners in the six temporal groups, whereas non-rhythmic anchor genes constituted the majority (72.9–77.1%; Fig. 6a). Rhythmic anchor genes exhibited distinct temporal interaction-degree profiles (Fig. 6b), while their expression phases were largely consistent with the phases of the dynamic interactions they participated in (Fig. 6c), indicating that gene expression oscillates in synchrony with chromatin loop formation.

**Fig. 6.**
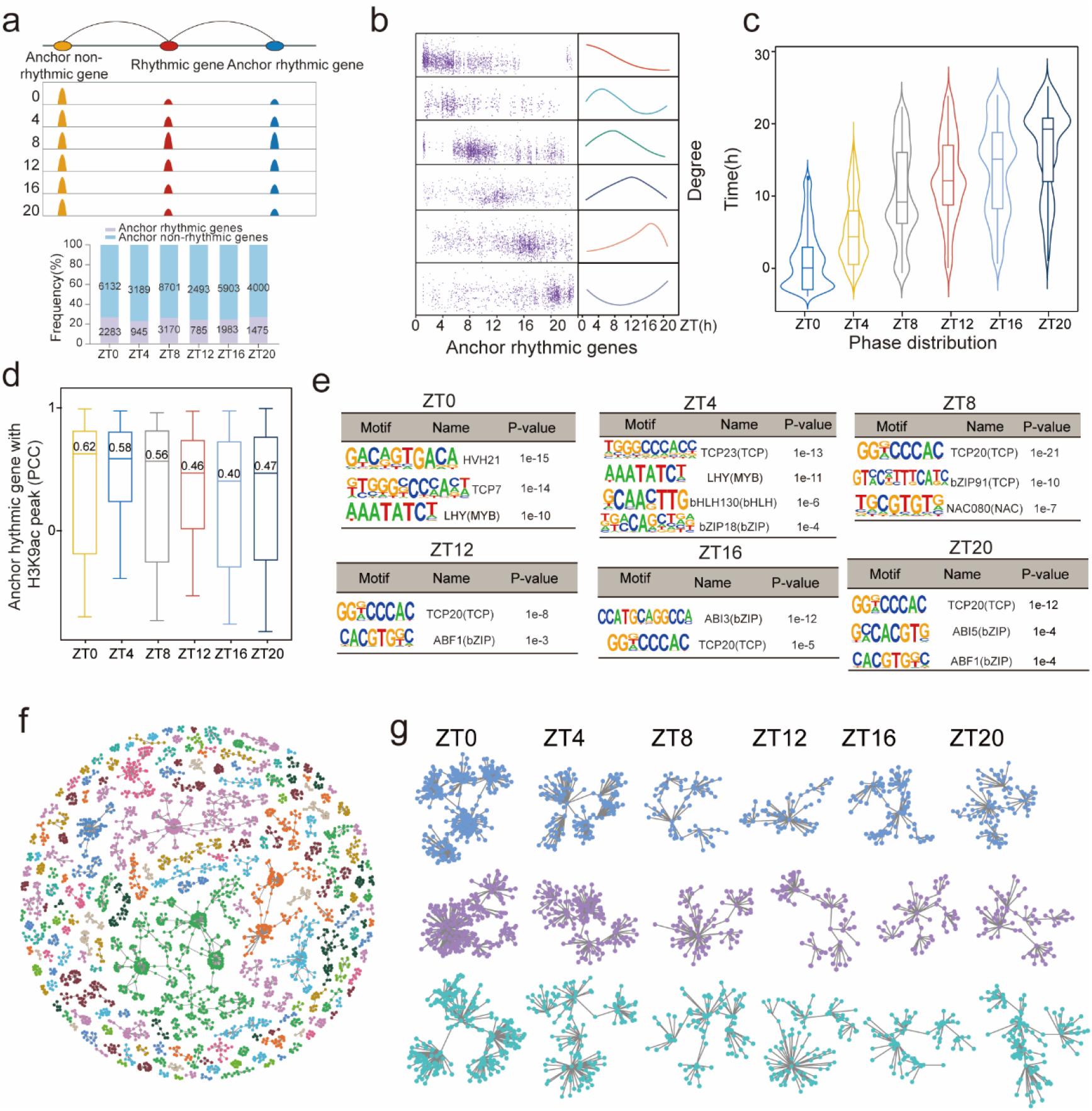
Dynamic local contacts and interaction networks involving rhythmic genes. a, Schematic identifying rhythmic anchor genes and distinguishing rhythmic and non-rhythmic genes participating in chromatin interactions (top). Stacked bars show their proportions in the ZT0, ZT4, ZT8, ZT12, ZT16 and ZT20 groups, with gene counts indicated. Rhythmic anchor-gene counts are 2,283, 945, 3,170, 785, 1,983 and 1,475, respectively. b, Temporal distributions of chromatin interaction degree for rhythmic anchor genes in the six groups (left) and the corresponding time-dependent profiles (right). c, Phase distributions of rhythmic anchor genes and their associated dynamic interactions across six temporal groups. d, Pearson correlation coefficients (PCCs) between rhythmic anchor-gene expression and associated H3K9ac peak signals across the six time points. Boxplots show the PCC distributions for each temporal group. e, Transcription factor binding motifs enriched in interaction-associated regions of the ZT0-ZT20 groups. Motif sequences, factor annotations and enrichment P values are shown. f, Network representation of interactions involving rhythmic genes. Nodes represent genes, edges represent detected interactions, and colors distinguish interaction groups. g, Temporal contact patterns of three representative gene groups across the six sampled time points. Colors distinguish groups; network layouts do not represent physical distances between loci.

To assess whether transcription and histone acetylation varied together at interaction anchors, we correlated the expression time series of rhythmic anchor genes with their associated H3K9ac peak signals. The PCC distributions had positive central values in all six groups, although individual gene–peak pairs varied substantially (Fig. 6d). Thus, transcriptional activity at interaction anchors was generally associated with H3K9ac dynamics over time. This correlation analysis measures temporal covariation; the relative timing of the two signals was examined separately using their estimated peak phases.

### Myb-and TCP-binding Motifs Are Enriched at Dynamic Chromatin Interaction Anchors

To uncover the regulatory factors governing dynamic loop rewiring, we analyzed DNA sequence motifs enriched at chromatin interaction anchors at each time point. LHY-like motifs were enriched at ZT0 and ZT4 (P = 1 × 10⁻¹⁰ and 1 × 10⁻¹¹, respectively), together with HVH21 and TCP7 at ZT0 and TCP23, bHLH130 and bZIP18 at ZT4. In the later groups, TCP20 motifs were enriched at ZT8, ZT12, ZT16 and ZT20, with ABF1-or ABI5-like motifs enriched in selected groups (Fig. 6e). These profiles identify candidate regulatory factors associated with interactions across the sampled cycle and reveal both shared and group-specific sequence enrichments.

Network representations provided examples of these interaction dynamics. Interactions involving rhythmic genes formed multiple connected gene groups (Fig. 6f), and three representative groups showed different patterns of detected contacts across the six time points (Fig. 6g). These examples illustrate temporal variation in chromatin interactions involving rhythmic genes within local networks.

Together, these analyses reveal dynamic chromatin interactions involving rhythmic genes and characterize their associated expression phases, histone acetylation and regulatory motifs within a largely conserved higher-order framework.

## Discussion

Our study demonstrates that diurnal changes in rice 3D genome organization occur within a largely stable, higher-order structural framework. While the majority of compartments and topologically associating domains (TADs) remained invariant across the 24-h cycle, selective genomic regions underwent dynamic compartment transitions, and local contacts fluctuated within conserved domains. Furthermore, a subset of cTADs spatially grouped rhythmic genes with synchronized expression phases. By integrating these multiscale structural features across time, our findings build upon previous snapshots of RNAPII-associated interactions and Hi-C analyses in rice [9,10], providing a continuous spatiotemporal view of the diurnal regulome.

Compartment switching captures regional variation within this predominantly stable landscape. In regions displaying prominent switching patterns, transitions into the active A compartment coincided with elevated RNAPII occupancy, increased H3K9ac enrichment, and diminished H3K9me2 levels. This aligns with previously reported links between active compartmentalization and open chromatin states in rice [5]. Notably, the pronounced clustering of oscillating chromatin compartments on chromosome 1 represents an intriguing phenomenon. Future investigations should evaluate how local gene density and chromatin landscape contribute to this chromosome-specific structural flexibility.

At the domain scale, conserved boundaries accompanied variable internal contacts, resembling observations in mouse liver [8]. In 35.8% of cTADs, rhythmic genes shared a phase category, indicating that a subset of domains spatially groups genes with similar expression phases. This organization is consistent with TADs providing a spatial framework for coordinated rhythmic transcription, although it does not establish a causal regulatory role. Because cTADs were also larger than ncTADs, comparisons controlling for domain size and gene content are needed to distinguish these structural properties from associations with rhythmic expression.

Beyond phase-synchronized chromatin domains, rhythmic genes also participated in dynamic chromatin interactions with both rhythmic and non-rhythmic partners. The rhythmic partners spanned broad expression-phase distributions, and representative network groups illustrated temporal changes in their contacts. These findings complement the shared expression phases observed within a subset of cTADs by showing that dynamic interactions also involve rhythmic genes with diverse phases. The network representations summarize pairwise contacts and do not establish simultaneous multiway contacts or physical clustering in individual nuclei.

Histone acetylation provides a common temporal context for the compartment and gene-network observations. H3K9ac and H3K27ac had the largest fractions of rhythmic features, whereas no rhythmic ATAC-seq features were detected under the applied criteria. This pattern is compatible with regulation on persistently accessible chromatin, as proposed in rice [10], and with links between histone acetylation and plant clock function [12], although detection limits preclude concluding that accessibility is invariant.

Motif enrichment identifies candidate transcription factors that may orchestrate these temporal interaction networks. LHY-like Myb-binding motifs were enriched at ZT0 and ZT4, whereas TCP20 motifs were detected in the ZT8–ZT20 groups. This temporal succession of enriched motifs suggests that stage-specific transcription factors may sequentially bind loop anchors to coordinate localized chromatin loop dynamics and maintain robust diurnal transcriptional cycles. Further study should focus on direct factor-occupancy profiling and targeted perturbations of candidate regulators or their binding elements to test their regulatory function.

Together, these findings support a model in which a globally conserved TAD framework maintains the spatial coordination of phase-coherent genes, whereas localized compartment transitions and dynamic chromatin loops actively rewire alongside transcriptional rhythms.

## Materials and methods

### Plant materials and growth conditions

Wild-type rice (*Oryza sativa* L. ssp. indica cv. Minghui 63; MH63) seedlings were grown in a phytotron under 12-h light/12-h dark cycles at 26 °C/24 °C and 70% relative humidity. Following synchronization, plants were transferred to constant light. Leaves were sampled every 4 h over 24 h, starting at ZT0, with two biological replicates per time point.

### ChIP-seq library preparation

Rice tissues were crosslinked with 1% formaldehyde and quenched with 0.2 M glycine. Approximately 0.5–1 g of tissue was ground in liquid nitrogen and suspended in buffer S (50 mM HEPES-KOH, 150 mM NaCl, 1 mM EDTA, 1% Triton X-100, 0.1% sodium deoxycholate, 1% SDS and protease inhibitors). Chromatin was fragmented to 200–600 bp at 4 °C using a Bioruptor (Diagenode). Lysates were centrifuged at 20,000 × g for 10 min at 4 °C, and the supernatant was collected. Anti-H3K9ac antibody (5–10 μL; Abcam, AB10812) was incubated with 200 μL of protein G magnetic bead suspension (Life Technologies, 10003D) for 1–6 h at 4 °C with rotation. Protein–DNA complexes were eluted in 100 μL of freshly prepared buffer (50 mM Tris-HCl, pH 7.5, 10 mM EDTA and 1% SDS). After reverse crosslinking, DNA was extracted with phenol:chloroform:isoamyl alcohol (Sigma-Aldrich, P3803), ethanol-precipitated and resuspended in TE buffer. Libraries were prepared using the NEBNext Ultra II DNA Library Prep Kit for Illumina (New England BioLabs, E7645). Fragments of 250–650 bp were selected using AMPure XP beads (Beckman, A63881) and sequenced on an Illumina HiSeq X Ten with 150-bp paired-end reads.

### RNA-seq library preparation

Total RNA was isolated from mature rice leaves using the RNeasy Plant Mini Kit (QIAGEN, 74904). Ribosomal RNA was depleted from 0.5–1.5 μg of total RNA using TruSeq Stranded Total RNA with Ribo-Zero Plant, following the manufacturer’s instructions. Libraries were sequenced on an Illumina HiSeq X Ten with 150-bp paired-end reads.

### BL-Hi-C library preparation

BL-Hi-C libraries were prepared as previously described [11], with modifications. Mature rice leaves were crosslinked with 1% formaldehyde (Sigma, F8775) for 10 min and 1.5 mM ethylene glycol bis(succinimidylsuccinate) (Thermo Fisher Scientific, 21565) for 30 min at room temperature. Crosslinking was quenched with 0.2 M glycine. Isolated nuclei were digested with MboI (NEB, R0147), followed by DNA end repair and A-tailing using T4 DNA polymerase (Promega, M421F) and Klenow enzyme (NEB, M0212L). Proximity ligation was performed in situ with a biotinylated bridge linker (forward: 5′-[5Phos]CGCGATATC/iBIOdT/TATCTGACT-3′; reverse: 5′-[5Phos]GTCAGATAAGATATCGCGT-3′). After reverse crosslinking, libraries were prepared using Tn5 transposase (VAHTS, TD501) and sequenced on an Illumina HiSeq X Ten with 150-bp paired-end reads (Annoroad, Beijing).

## Analysis methods

### RNA-seq data processing

RNA-seq reads were assessed with FastQC v0.11.9 and trimmed using Trimmomatic v0.36 [13] with default parameters. Filtered reads were aligned to the MH63 genome using HISAT2 v2.1.0 [14]. Gene counts were obtained with HTSeq v0.11.2 [15] and normalized using DESeq2 in R [16].

### ChIP-seq data processing

ChIP-seq reads were assessed with FastQC, filtered with Trimmomatic and aligned to the MH63 genome using BWA-MEM v0.7.17 [17] with default parameters. PCR duplicates were removed using SAMtools v1.9 rmdup [18], and reads with mapping quality >30 were retained. Peaks were called using MACS2 v2.1.1 [19] with -f BAM -g 3.6e+8 and an input control specified by -c. Samples were required to have relative and normalized strand cross-correlation coefficients >0.8 and >1.05, respectively. Enrichment was assessed using the fraction of reads in peaks (FRiP). Normalized bigWig tracks were generated using deepTools v3.2.1 bamCoverage [20] and visualized in IGV v2.5.0 [21].

### BL-Hi-C data processing

BL-Hi-C reads were processed with the ChIA-PET pipeline [22] for linker filtering, alignment and interaction calling. Linker sequences were specified as -A GATCACGCGATATCTTATCTGACTGATC and -B GATCAGTCAGATAAGATATCGCGTGATC. Interactions with a false discovery rate (FDR) <0.05 and ≥4 PETs were retained.

### BL-Hi-C contact map construction and visualization

Intrachromosomal inter-ligation PETs spanning >8 kb were retained for contact maps. PET files were converted to .hic format using Juicer tools v1.8.9 [23] with default parameters and visualized in Juicebox v1.11.08 [24].

### Time-series analysis of rhythmicity

Normalized RNA-seq and ChIP-seq time series were analyzed using BIO_CYCLE [25] with a candidate period range of 20–28 h and otherwise default parameters. Features with P < 0.05 were classified as rhythmic, and their estimated periods, phases and amplitudes were retained.

### A/B compartment assignment and TAD identification

A/B compartments were assigned from PC1 of chromosome-wise contact correlation matrices using the Juicer eigenvector tool [23]. Regions that changed compartment identity across the six time points were designated OCCs; unchanged regions were classified as stable. Matrices were converted from .hic to .cool format using hicConvertFormat, and TADs were identified with hicFindTADs in HiCExplorer v3.5.1 [26].

### Chromatin interaction network visualization

BL-Hi-C interaction networks were visualized in Gephi v0.9.3 [27], using node color, node size and edge color to distinguish displayed features. For the genome-wide ZT0 network, 20% of interactions were randomly sampled for visualization.

### Transcription factor motif enrichment

Motif enrichment was analyzed separately for each temporal group using HOMER [28] in regions associated with interactions involving rhythmic genes. Plant motifs were selected with the reported parameter -set plants.

